# Phenotypic Bioactivity Prediction as Open-set Biological Assay Querying

**DOI:** 10.64898/2026.02.28.708711

**Authors:** Yuze Sun, Xiaoman Zhang, Qiaoyu Zheng, Hanzheng Li, Jianming Zhang, Liang Hong, Yanfeng Wang, Ya Zhang, Weidi Xie

**Affiliations:** School of Artificial Intelligence, Shanghai Jiao Tong University, Shanghai, China; Shanghai Artificial Intelligence Laboratory, Shanghai, China; Department of Biomedical Informatics, Harvard Medical School, Boston, MA, USA; Institute of Natural Sciences, Shanghai Jiao Tong University, Shanghai, China; Institute of Translational Medicine, Zhangjiang Institute for Advanced Study, Shanghai Jiao Tong University, Shanghai, China; School of Physics and Astronomy & Institute of Natural Sciences, Shanghai Jiao Tong University, Shanghai, China; Shanghai National Centre for Applied Mathematics (SJTU Center), MOE-LSC, Shanghai Jiao Tong University, Shanghai, China; Zhangjiang Institute for Advanced Study, Shanghai Jiao Tong University, Shanghai, China

## Abstract

The traditional drug discovery pipeline is severely bottlenecked by the need to design and execute bespoke biological assays for every new target and compound—a process that is both time-consuming and prohibitively expensive. While machine learning has accelerated virtual screening, current models remain confined to “closed-set” paradigms, unable to generalize to entirely novel biological assays without target-specific experimental data. Here, we present **OpenPheno**, a groundbreaking multimodal foundation model that fundamentally redefines bioactivity prediction as an open-set, visual-language question-answering (QA) task. By integrating chemical structures (SMILES), universal phenotypic profiles (Cell Painting images), and natural language descriptions of biological assays, OpenPheno unlocks the highly coveted **“profile once, predict many” paradigm**. Instead of conducting countless target-specific wet-lab experiments, researchers only need to capture a single, low-cost Cell Painting image of a novel compound. OpenPheno then evaluates this universal phenotypic “fingerprint” against the text-based description of any unseen assay, predicting bioactivity in a zero-shot manner. On 54 entirely unseen assays, it achieves strong zero-shot performance (mean AUROC 0.75), exceeding supervised baselines trained with full labeled data, and few-shot adaptation further improves predictions. In the most stringent setting where both compounds and assays are novel, OpenPheno maintains robust generalization (mean AUROC 0.66), opening up a new paradigm for a highly scalable, cost-effective, and universal engine for next-generation drug discovery.

## 1 Introduction

Drug discovery is limited by the cost and latency of evaluating therapeutic hypotheses in large-scale screenings [1]. Unbiased profiling technologies, including high-content imaging (e.g., Cell Painting [2]) and transcriptomic perturbation assays (e.g., L1000 [3]), address this bottleneck by capturing broad, system-level cellular responses to chemical perturbations. These rich representations allow researchers to identify bioactive compounds without predefining molecular targets [4, 5]. A central computational challenge is “phenotypic bioactivity prediction”: predicting whether a compound will exhibit activity in a specific assay by integrating chemical structures with phenotypic profiles.

However, current computational approaches are limited by a rigid formulation of this problem. Existing methods treat bioactivity prediction as closed-set multi-label classification, in which the set of assays is fixed at training time and the model is trained to predict outcomes only for this pre-determined panel [6–10]. This fails to reflect the dynamic reality of drug discovery, where biological questions evolve, and new assays are continuously developed. Consequently, identifying active compounds for a novel assay currently requires re-screening or re-training models from scratch. A truly scalable framework must generalize to assays never seen during training—a capability that closed-set models fundamentally lack.

Here, we present **OpenPheno, a multimodal framework that reformulates bioactivity prediction from closed-set classification to open-set biological prediction (Fig. 1a)**. By integrating Cell Painting images, chemical structures, and natural-language assay descriptions, OpenPheno learns to predict whether a compound exhibits a queried biological assay, enabling generalization to both unseen assays and novel compounds without retraining. **Our approach employs a two-stage training strategy:** first, a large-scale multimodal pretraining that aligns visual representations with chemical structures while enforcing consistency across experimental replicates through self-supervised learning; and second, an assay query network (AQN) that enables dynamic predictions by fusing assay descriptions with compound features.

**Figure 1.**
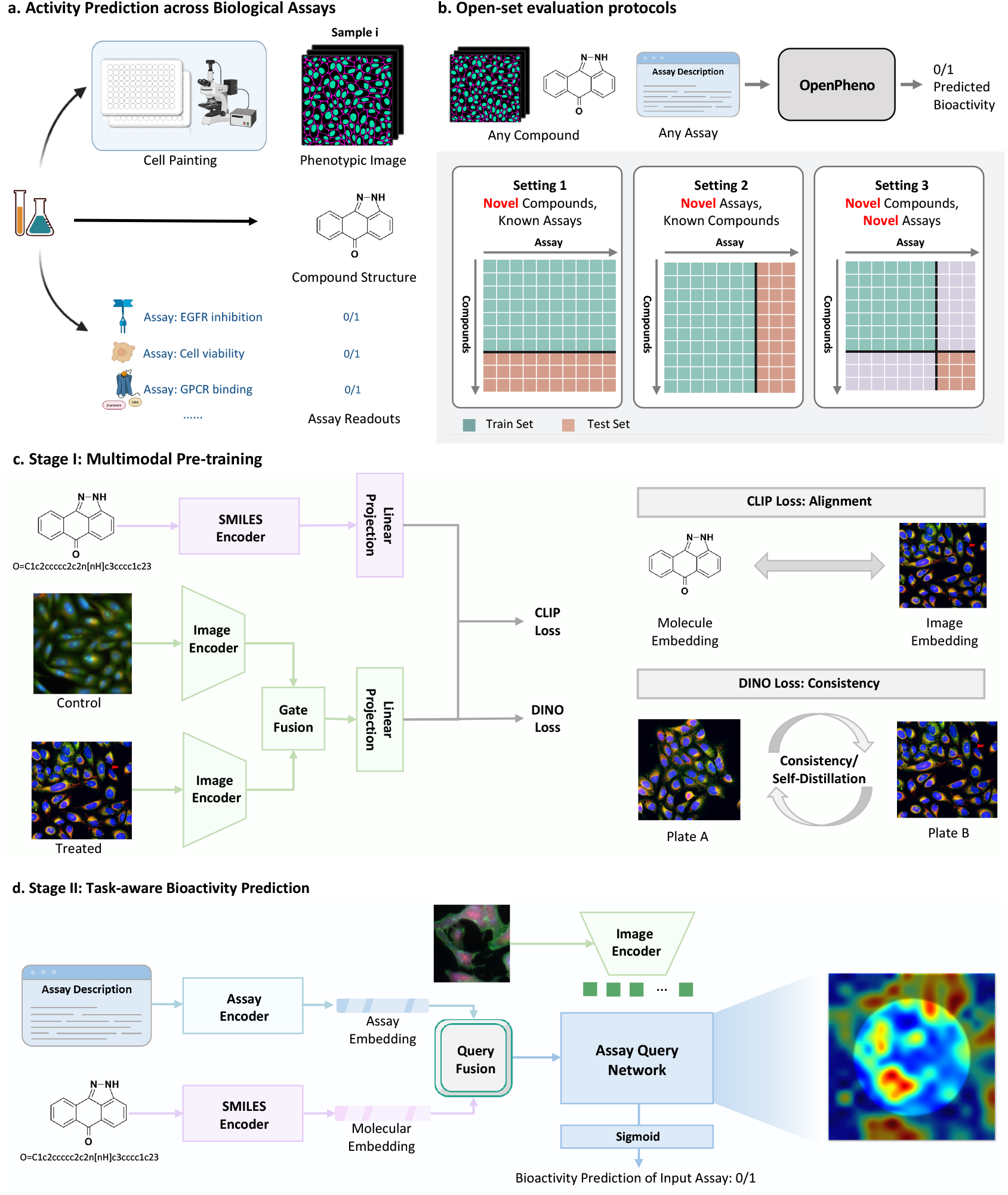
Overview of OpenPheno for open-set bioactivity prediction. **a**. Task formulation: a compound is represented by its Cell Painting image and chemical structure; given a natural-language assay description as query, bioactivity is predicted as a binary outcome. **b**. Three evaluation protocols of increasing difficulty. Setting 1: novel compounds, known assays. Setting 2: known compounds, novel assays. Setting 3: both novel. **c**. Stage I multimodal pretraining combines cross-modal contrastive alignment (CLIP loss) with cross-plate self-supervised consistency (DINO loss). **d**. Stage II assay-aware prediction: the Assay Query Network fuses compound and assay description embeddings into a dynamic query that attends over image patch features, enabling zero-shot generalization to unseen assays.

We evaluated OpenPheno on two established benchmarks: Broad-270 [9] (16,170 compounds, 270 assays) and ChEMBL-209 [7] (10,574 compounds, 209 assays), across all three generalization settings. **The first setting** resembles the existing literature, where novel compounds are tested against known assays, OpenPheno outperformed all existing models on both benchmarks (mean AUROC **0.76** on ChEMBL-209 and **0.71** on Broad-270), confirming that open-set flexibility does not sacrifice closed-set predictive performance. In the **second setting**, where known compounds are evaluated against 54 assays entirely withheld from training, OpenPheno achieved accurate zero-shot predictions using only assay descriptions (mean AUROC **0.75**), and few-shot adaptation with limited labeled data further improved performance. In the **third and most challenging setting**, where both compounds and assays are unseen during training, OpenPheno maintained strong zero-shot generalization (mean AUROC **0.66**). Notably, across both open-set settings, OpenPheno’s zero-shot performance surpassed that of all baseline models even when those baselines were fine-tuned on target assay labels (best baseline AUROC **0.64**). These results demonstrate that recasting bioactivity prediction as open-set biological querying enables generalization across the expanding landscape of biological assays without sacrificing performance on established benchmarks.

## 2 Results

We present **OpenPheno**, a multimodal framework that reformulates bioactivity prediction as open-set biological querying. The framework employs two-stage training: **Stage I** pretrains a visual encoder to align phenotypic and molecular representations while enforcing cross-replicate consistency; **Stage II** introduces an assay query network (AQN) that enables predictions by conditioning on assay descriptions, enabling zero-shot inference. We evaluate OpenPheno across three settings that test increasingly challenging generalization scenarios: (i) unseen compound generalization, where novel compounds are tested against known assays; (ii) unseen assay generalization, where known compounds are evaluated against entirely unseen biological assays; and (iii) unseen compound and assay generalization, where both compounds and assays are novel simultaneously.

### 2.1 Benchmark Description

#### Datasets

All phenotypic images are derived from the Cell Painting assay [2], a high-content microscopy protocol that captures broad morphological responses to chemical perturbations. Bioactivity ground-truth labels are drawn from two large-scale screening benchmarks:

i. **Broad-270** [9], comprising 16,170 compounds tested across 270 assays spanning cell-based, biochemical, bacterial, fungal, and yeast. To facilitate binary classification, continuous assay readouts were binarized via statistical thresholding following double-sigmoid normalization [11]. This dataset serves as the primary benchmark for all three evaluation settings, as described in the evaluation settings;
ii. **ChEMBL-209** [7], comprising 10,574 compounds across 209 assays reflecting large-scale screening conditions. Here, binary activity labels were assigned based on standardized pChEMBL value thresholds or explicit activity annotations. This dataset is used for unseen compound generalization (Setting 1) to assess performance across an independent benchmark.

#### Evaluation settings

This paper considers three protocols (Fig. 1b), as detailed below:

- **Setting 1: generalization to unseen compounds**. This setting follows the standard protocol established in prior work [7, 9], where all assays are observed during training, but test compounds are structurally distinct from training ones (scaffold-based partitioning). On Broad-270, compounds are partitioned by Bemis-Murcko scaffolds [12], which represent the core ring systems and linkers of each molecule; all compounds sharing a scaffold are assigned exclusively to either the training or test set, ensuring that the model is evaluated on structurally novel chemical matter. On ChEMBL-209, compounds are partitioned following the original benchmark protocol [7].
- **Setting 2: generalization to unseen assays**. This setting assesses the model’s ability to predict activity for biological assays not encountered during training. 54 of the 270 assays in Broad-270 are entirely withheld from training. We evaluate performance in two modes: zero-shot inference, where predictions rely solely on assay descriptions and all compounds associated with the held-out assays serve as the test set; and few-shot adaptation, where the compounds within each held-out assay are further partitioned by molecular scaffold into support and query sets, such that fine-tuning and evaluation are performed on structurally distinct compounds.
- **Setting 3: generalization to unseen compounds and assays**. This setting denotes the most rigorous evaluation, requiring simultaneous generalization to both novel chemical compounds and novel biological tasks. Test compounds are scaffold-separated from the training set, and test assays are entirely unseen. This setting effectively measures the capacity of OpenPheno to address new biological questions regarding novel chemical entities, the scenario most critical for real-world drug discovery.

### 2.2 OpenPheno Generalizes to Unseen Compounds

We first evaluate OpenPheno under the standard compound-split protocol on both benchmarks (Fig. 2a), where all assays are observed during training, and test compounds are structurally distinct from training compounds via scaffold-based partitioning. For comparison, we adopt state-of-the-art bioactivity prediction methods spanning the following two categories: (i) **Supervised models** trained end-to-end on raw microscopy images. On ChEMBL-209, this category includes ResNet-50 [13], DenseNet-121 [14], GapNet [7], MIL-Net [15], M-CNN [16], and SC-CNN [7], for which we report original published results [7]. On Broad-270, we retrain ResNet-50, the best-performing architecture in this family, under identical data splits for fair comparison. (ii) **Linear probing** on frozen representations. This category encompasses two subtypes. **Feature-based approaches:** on Broad-270, LateFusion [9] trains separate multi-task feedforward networks on pre-computed chemical structure (CS), Cell Painting morphology (MO), and gene expression (GE) features before aggregating prediction scores; on ChEMBL-209, we evaluate a feedforward network trained on CellProfiler morphological features [6]. **Contrastive pretraining:** CLOOME [10] pre-trains a visual encoder via contrastive learning to align microscopy images with chemical structures, followed by linear probing for bioactivity prediction. We additionally report linear probes on our own frozen encoders (Vision Encoder and SMILES Encoder from Stage I pretraining) to isolate the contribution of the Assay Query Network.

**Figure 2.**
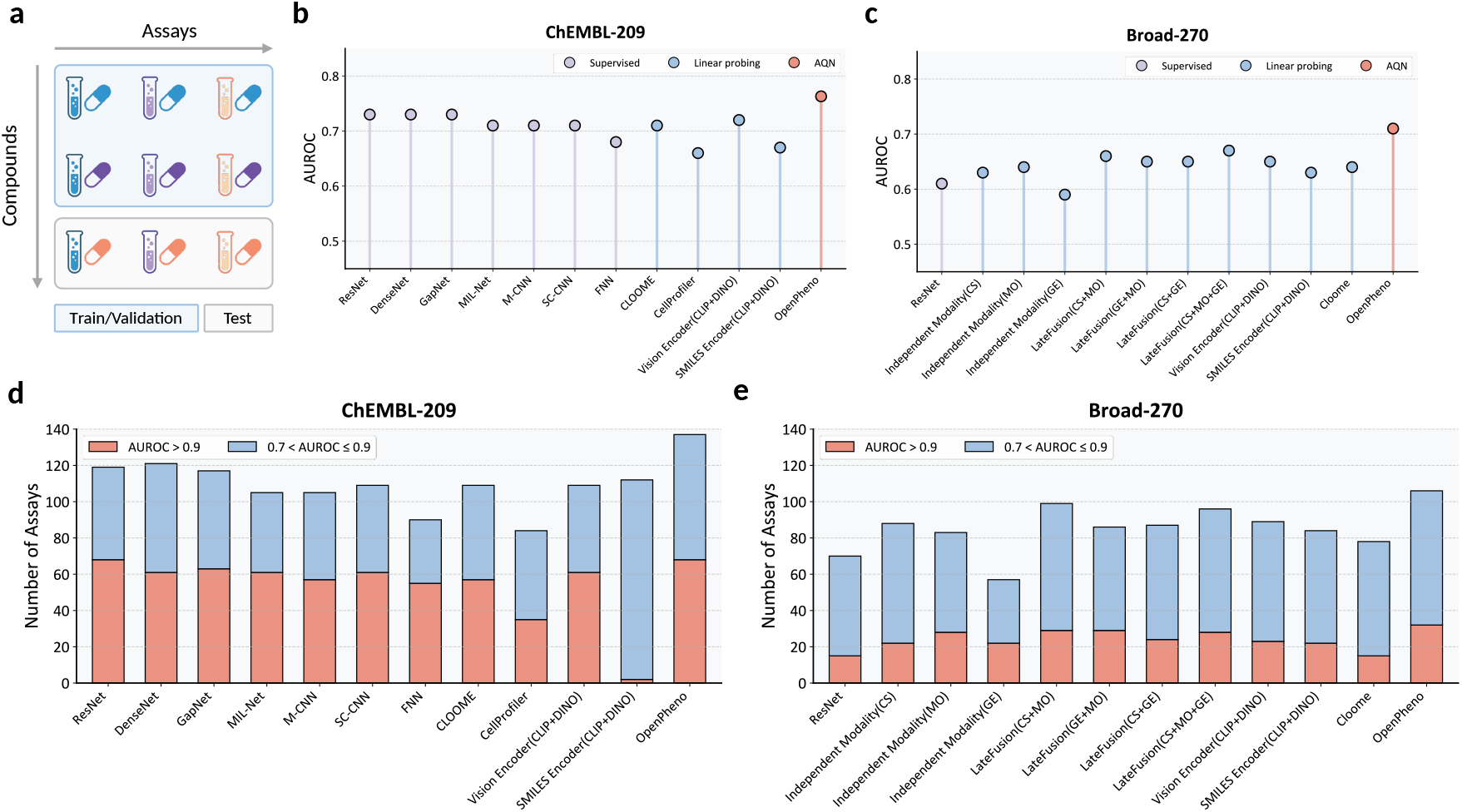
Closed-set bioactivity prediction on standard benchmarks. **a**. Evaluation protocol: all assays are observed during training; test compounds are structurally distinct from training compounds (scaffold-based split). **b**,**c**. Mean AUROC across all assays on ChEMBL-209 (**b**) and Broad-270 (**c**). Methods are grouped by type: supervised models trained end-to-end on raw pixels (grey), linear probing on frozen representations (blue), and OpenPheno with the Assay Query Network (red). **d**,**e**. Number of assays exceeding AUROC thresholds of 0.9 (high-confidence, red) and 0.7 (useful for screening, blue) on ChEMBL-209 (**d**) and Broad-270 (**e**). OpenPheno achieves the highest mean AUROC on both benchmarks and the largest number of assays above AUROC 0.7, despite using an open-set architecture not specifically optimized for closed-set prediction.

#### Performance on Broad-270

On Broad-270 (Fig. 2c,d; Table S1), OpenPheno achieved a mean AUROC of **0.71**, surpassing the best late fusion baseline that combines all three data modalities (LateFusion CS+MO+GE: 0.67) and single-modality approaches (CS: 0.63; MO: 0.64; GE: 0.59). At the AUROC > 0.7 threshold, OpenPheno predicted 106 assays compared to 99 for LateFusion (CS+MO) and 96 for LateFusion (CS+MO+GE). Notably, OpenPheno achieves this using only phenotypic images and chemical structures, without gene expression data, yet outperforms the three-modality fusion, indicating that learned cross-modal attention integrates complementary signals more effectively than post hoc score aggregation.

#### Performance on ChEMBL-209

On ChEMBL-209 (Fig. 2b,e; Table S2), OpenPheno achieved a mean AUROC of **0.76**, outperforming all supervised baselines trained end-to-end on raw images, including ResNet-50 (0.73) and DenseNet-121 (0.73), contrastive pretraining approaches (CLOOME: 0.71), and feature-based methods (CellProfiler: 0.66). At the AUROC > 0.7 threshold, OpenPheno predicted 137 assays, exceeding all baselines. The consistent improvements across both benchmarks confirm that conditioning predictions on assay semantics provides complementary benefits even when all assays are observed during training.

### 2.3 OpenPheno Generalizes to Unseen Assays

We then evaluate OpenPheno under the assay-split protocol on Broad-270 (Fig. 3a), where 54 of the 270 assays are entirely withheld from training, and the model must predict compound activity using only natural-language assay descriptions. Since existing closed-set methods require all assays to be defined at training time, they cannot operate in this setting. We evaluate performance in two modes: zero-shot inference, where no labeled examples are provided for the target assay; and few-shot adaptation, where increasing fractions of labeled data are used for fine-tuning.

**Figure 3.**
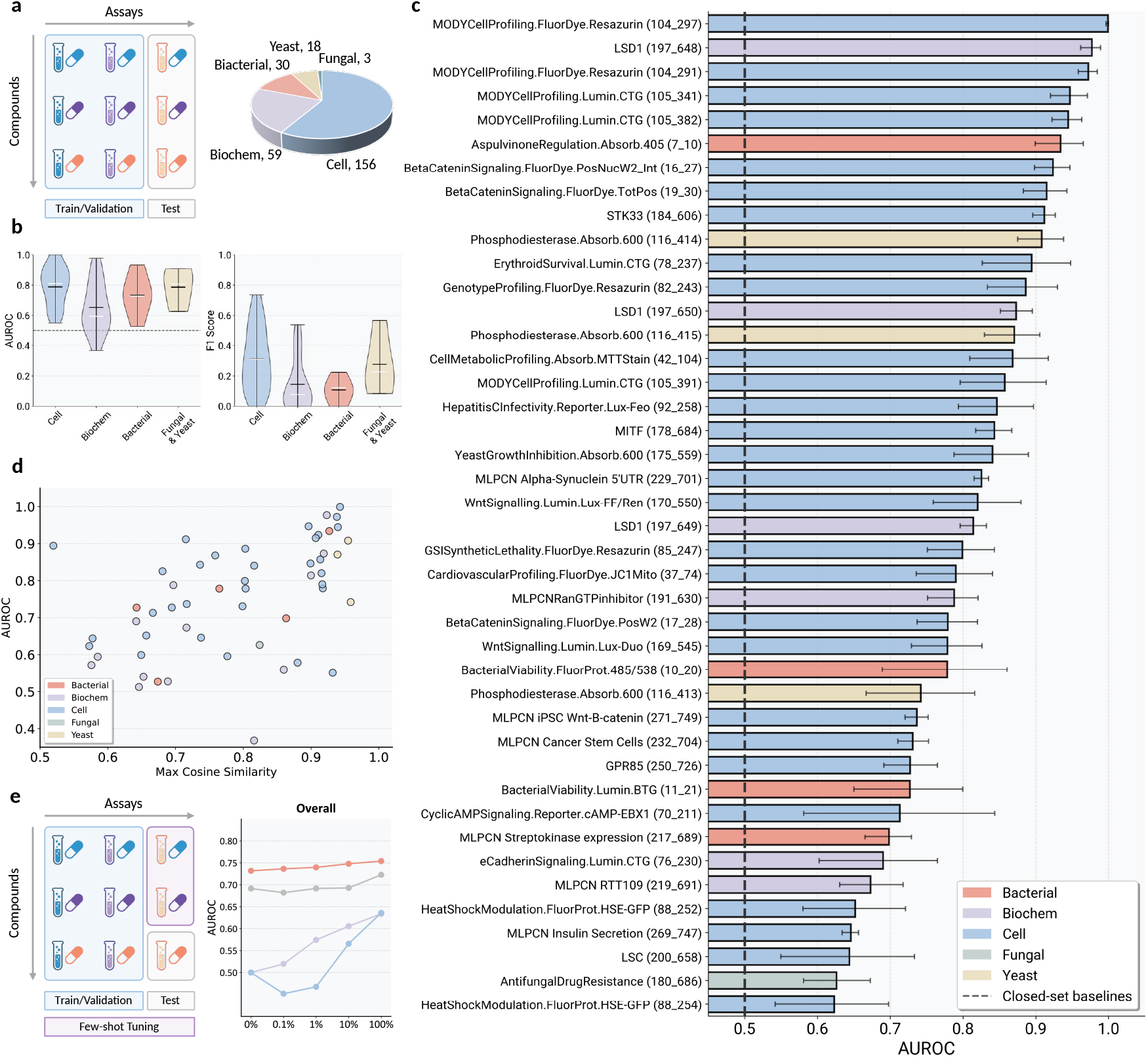
OpenPheno enables zero-shot bioactivity prediction across diverse unseen assays. **a**. Assay-split evaluation protocol with the distribution of assay categories. **b**. Zero-shot performance stratified by assay category, reported as AUROC (left) and F1 score (right). **c**. Per-assay zero-shot AUROC for all 54 held-out assays, ranked by performance. **d**. Relationship between zero-shot AUROC and maximum cosine similarity of each test assay description to the training set. Performance correlates positively with semantic similarity.

#### Overall performance

As shown in Figure 3, on the 54 held-out assays from Broad-270, OpenPheno achieved a mean AUROC of 0.75 (95% CI: 0.70–0.80; Table S3) in zero-shot mode, using only natural-language assay descriptions to predict compound activity without any labeled examples for the target assay.

#### Performance across assay categories

We stratified zero-shot performance by assay category (Fig. 3b,c). Cell-based assays achieved the highest zero-shot performance (mean AUROC 0.79, 95% CI: 0.74–0.83), consistent with the direct correspondence between Cell Painting morphological readouts and cell-based experimental questions. Fungal and yeast assays also showed strong generalization (mean AUROC 0.79, 95% CI: 0.74–0.83), though with wider confidence intervals reflecting the smaller sample size (*n* = 3). Bacterial assays maintained robust performance (mean AUROC 0.73, 95% CI: 0.68–0.78), indicating that the learned representations capture biological principles that transfer across organisms despite training predominantly on mammalian cell phenotypes. Biochemical assays proved more challenging (mean AUROC 0.65, 95% CI: 0.61–0.70), as expected given that purified molecular interactions lack direct morphological correlates in Cell Painting images. Nevertheless, even for biochemical assays, OpenPheno substantially exceeded the random baseline of 0.5, suggesting that compound-induced phenotypic signatures retain information about target-level biochemical activity.

#### Relationship between semantic similarity and prediction accuracy

To characterize the conditions under which zero-shot prediction succeeds, we examined the relationship between test assay performance and semantic similarity to the training set (Fig. 3d). For each test assay, we computed the maximum cosine similarity between its embedding and all training assays, *i*.*e*.,computed by encoding the assay description with BioLord [17]. Test assay AUROC was positively correlated with this similarity measure (Spearman *ρ* = 0.551, *p <* 0.001), confirming that OpenPheno leverages semantic relationships between assay descriptions to transfer knowledge from related training contexts. Notably, several assays with moderate semantic similarity (cosine *<* 0.7) still achieved AUROC above 0.8, suggesting that the model generalizes beyond simple description matching by exploiting shared phenotypic signatures across biologically related experiments.

#### OpenPheno achieves data-efficient adaptation to novel assays

While zero-shot inference provides strong baseline performance, practical drug discovery often allows for a small pilot screen. We evaluated few-shot adaptation by fine-tuning OpenPheno on increasing fractions (ranging from 0.1% to 100%) of labeled data from the 54 unseen assays (Fig. 3e). As shown in Fig. 3e and Table S4, OpenPheno exhibits stable performance across all data regimes. Even at 0.1% labeled data, OpenPheno achieves a mean AUROC of (95% CI: 0.66–0.81), marginally improving over its zero-shot performance (0.73) and already exceeding all baselines at full data utilization (SMILES Encoder: 0.63 at 100%; Vision Encoder: 0.64 at 100%). With 10% of the data, performance reaches 0.75 (95% CI: 0.68–0.82), and at full supervision the model achieves (95% CI: 0.67–0.83), indicating that the pretrained representations already capture the majority of predictive signal. We also examined how few-shot adaptation varies across assay categories(Table S4). Across all categories, baseline learning curves plateau below OpenPheno’s zero-shot level, confirming that multimodal pretraining provides a fundamentally stronger initialization than task-specific training from scratch.

### 2.4 OpenPheno Generalizes to Both Unseen Compounds and Unseen Assays

Setting 3 represents the most demanding evaluation scenario, requiring simultaneous generalization to structurally novel compounds and entirely unseen biological assays. Unlike Settings 1 and 2, which isolate each axis of generalization independently, **this setting directly assesses the capacity of OpenPheno to address new biological questions about new chemical entities, the scenario most critical to real-world drug discovery workflows, where candidate compounds and the assays used to characterize them are both unknown at the time of model training**.

#### Overall performance

On the 54 held-out assays from Broad-270 (Fig. 4a,c), OpenPheno achieved a mean AUROC of 0.66 in zero-shot mode, despite having access neither to labeled examples for the target assays nor to any compounds sharing a molecular scaffold with the test set. This performance substantially exceeds the best closed-set baseline trained with full supervision on the target assays (AUROC 0.64), confirming that OpenPheno’s multimodal representations and assay-conditioned query mechanism retain discriminative power even when both generalization challenges are imposed jointly.

**Figure 4.**
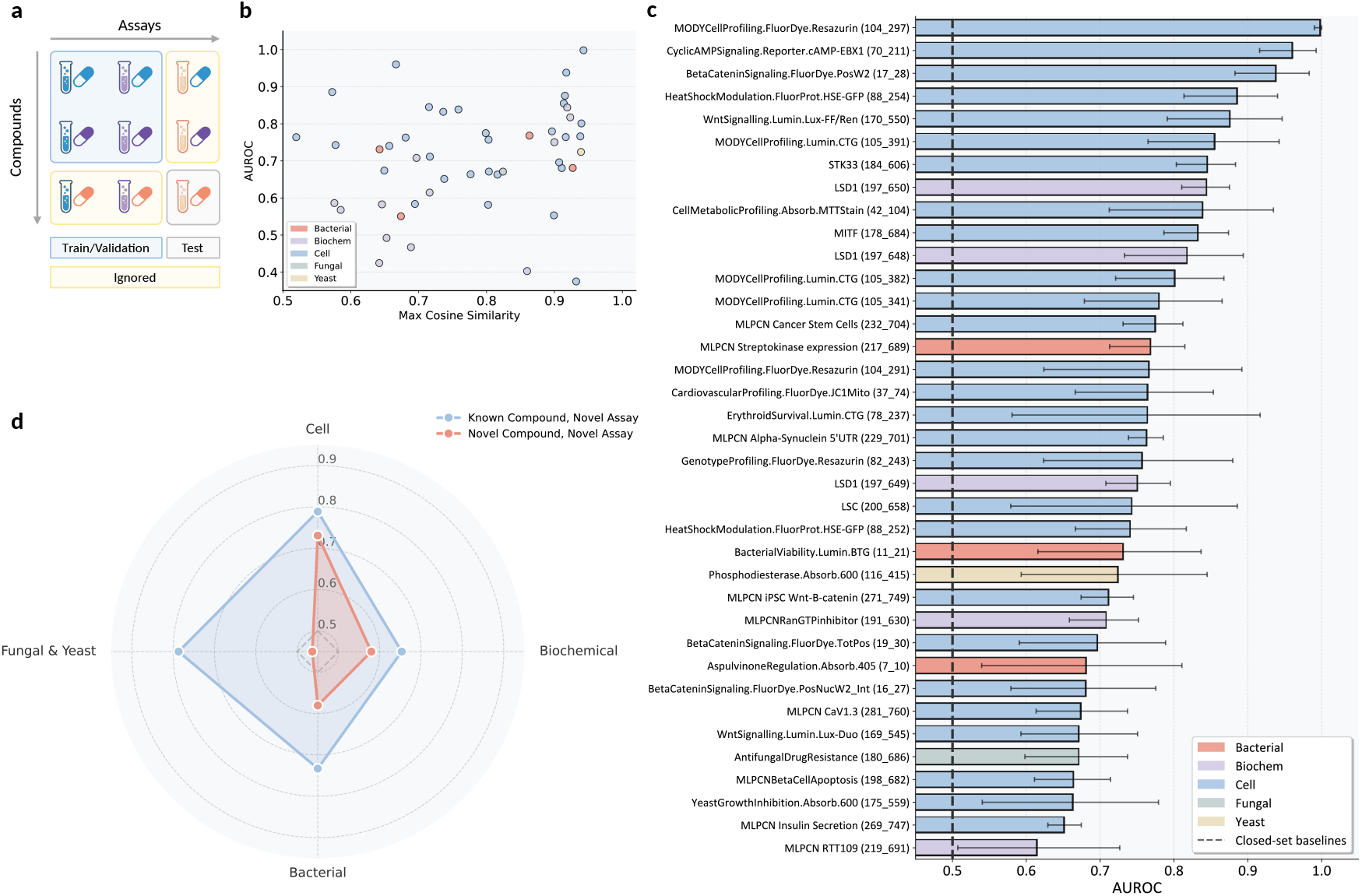
OpenPheno generalizes to the fully open-set setting (novel compounds and novel assays). **a**. Evaluation protocol for Setting 3: the most rigorous scenario, where test compounds are structurally distinct (scaffold-split) from training data and are evaluated on biological assays never seen during training. **b**. Relationship between zero-shot AUROC and maximum cosine similarity of each test assay description to the training set. Performance correlates positively with semantic similarity. **c**. Per-assay zero-shot AUROC for all 54 held-out assays, ranked by performance. **d**. Radar chart comparing the mean zero-shot AUROC across four biological categories between Setting 2 and Setting 3.

#### Relationship between performance and semantic similarity

As in Setting 2, zero-shot AUROC in Setting 3 correlates positively with the maximum cosine similarity between each held-out assay description and the training assay descriptions (Fig. 4b), confirming that semantic knowledge transfer through assay descriptions operates independently of whether the test compounds were represented during training. The performance degradation from Setting 2 to Setting 3 is therefore attributable primarily to compound structural novelty rather than to a failure of assay-level generalization, suggesting that improving the robustness of phenotypic representations to chemical novelty represents the primary avenue for further gains in this setting.

#### Performance across assay categories

The radar chart in Fig. 4d directly compares category-level zero-shot performance between Setting 2 (known compounds, novel assays) and Setting 3 (novel compounds, novel assays). For cell-based assays, performance remains closely matched across the two settings, indicating that when assays directly interrogate morphological phenotypes, compound structural novelty has minimal impact on predictive accuracy. In contrast, biochemical assays show the largest gap between settings, suggesting that for purified molecular interactions lacking direct morphological correlates, the model relies more heavily on having encountered chemically similar compounds during training. Together, these results demonstrate that the robustness of OpenPheno to compound novelty is primarily determined by the degree to which the target assay is grounded in cellular phenotypes, and that phenotypic imaging provides a modality-level generalization advantage that is largely independent of chemical structure.

### 2.5 Multimodal Pretraining Establishes Robust Feature Foundations

#### Ablation of pretraining objectives

We compared the full dual-objective pretraining (CLIP+DINO) against single objectives and alternative strategies on the unseen assay generalization task(Table S6; Fig. 5a). Training from scratch yielded a mean AUROC of 0.72, establishing the reference baseline. MAE [18] underperformed even this baseline (0.70), likely because pixel-level reconstruction encourages encoding of plate-specific artifacts that are uninformative for bioactivity prediction. SigLIP [19] achieved a mean AUROC of 0.73, marginally below the standard InfoNCE-based CLIP objective (mean AUROC 0.74), suggesting that softmax-normalized contrastive learning provides more effective cross-modal alignment in this domain. Notably, the two retained objectives exhibit complementary strengths across assay categories. DINO achieved the highest single-objective performance on cell-based assays (0.80) but the lowest on biochemical assays (0.59), indicating that cross-plate self-distillation captures morphological consistency effectively but lacks the molecular context needed for non-morphological screens. CLIP showed the opposite pattern, with stronger biochemical performance (0.65) driven by chemical structure grounding. The full model (CLIP+DINO, 0.75) exceeded either objective alone across all categories, confirming a synergistic effect: DINO suppresses plate-level batch effects while CLIP injects molecular semantics, together providing a comprehensive foundation for the Assay Query Network.

**Figure 5.**
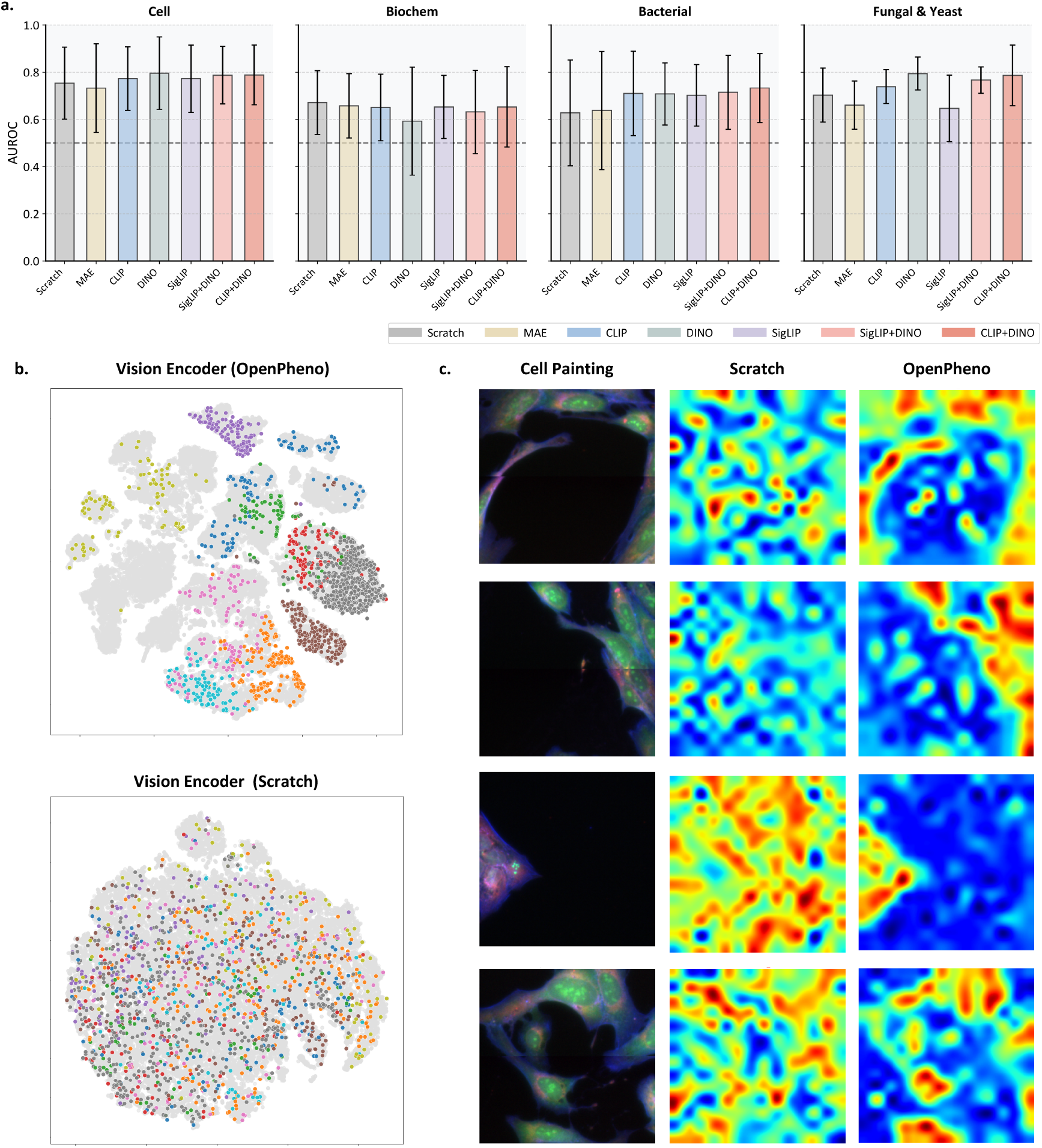
Multimodal pretraining establishes robust representations for bioactivity prediction. **a**. Ablation of Stage I pretraining objectives on the unseen assay generalization task (54 held-out assays), stratified by assay category. **b**. t-SNE visualization of compound embeddings from the OpenPheno vision encoder (top) and a vision encoder without pretraining (bottom). Grey points represent the full test set distribution, while colored points highlight 10 representative clusters corresponding to distinct scaffold classes. **c**. Cross-attention maps from the Assay Query Network before (middle column) and after (right column) Stage II training, alongside the original Cell Painting images (left column), visualized as pseudo-color RGB composites constructed from the ERSyto, ERSytoBleed, and Ph_golgi channels.

#### t-SNE visualization of embedding space

We visualized the learned representations using t-SNE [20] to assess how pretraining shapes the feature landscape (Fig. 5b). Without pretraining, compound embeddings appear scattered with minimal organization. In contrast, OpenPheno produces a structured latent space in which compounds sharing molecular scaffolds form coherent clusters, indicating that the dual-objective pretraining projects high-dimensional microscopy data into a biologically meaningful manifold that facilitates downstream assay querying.

#### Attention maps confirm biologically grounded predictions

To verify that the Assay Query Network leverages biologically relevant features, we visualized its cross-attention maps before and after Stage II training (Fig. 5c). Before training, attention is diffuse and distributed across background regions. After training, the model consistently attends to subcellular structures, including nuclei, perinuclear regions, and cytoplasmic organelles, while suppressing background noise. This provides evidence that OpenPheno grounds its predictions in biologically relevant cellular phenotypes rather than imaging artifacts.

## 3 Discussion

### Open-set querying as a new paradigm for bioactivity prediction

Current computational approaches treat bioactivity prediction as closed-set classification, restricting inference to assays seen during training [6,7,9, 10]. This is mismatched with drug discovery, where biological questions evolve and new assays are continuously developed [5]. OpenPheno addresses this gap by recasting each assay as a natural-language query rather than a fixed output index, enabling the model to interpret biological questions never encountered during training. The positive correlation between assay description similarity and zero-shot accuracy (Fig. 3d) confirms that OpenPheno transfers knowledge through shared biological semantics, while strong performance on assays with only moderate semantic similarity suggests generalization beyond simple description matching. This distinguishes OpenPheno from prior strategies: ActFound [21] addresses assay incompatibility via pairwise meta-learning but still requires labeled examples at inference; CLOOME [10] aligns images with structures via contrastive learning but lacks assay semantics, limiting it to the closed-set regime; and TxGNN [22] achieves zero-shot drug repurposing over knowledge graphs at the disease-indication level rather than individual assays. To our knowledge, OpenPheno is the first framework to enable zero-shot bioactivity prediction conditioned on assay descriptions at the granularity of individual biological experiments.

### Multimodal pretraining enables data-efficient deployment of new assays

OpenPheno’s zero-shot performance on unseen assays exceeds supervised baselines trained on the full labeled dataset, and few-shot adaptation with only 0.1% of labeled data improves further, implying that deploying a new assay could shift from large-scale screening to a small pilot experiment. The dual-objective pretraining strategy underlies this efficiency: image-wise self-supervised learning suppresses plate-level batch effects critical for cell-based assays, while contrastive learning for alignment injects the molecular context needed for biochemical screens. In ablation studies, combining both objectives outperformed using CLIP or DINO alone across cell-based, bacterial, and biochemical assay categories (Fig. 5), confirming that the two objectives capture complementary features. In contrast, generic self-supervised strategies such as masked autoencoders, which lack these biology-specific inductive biases, degraded performance relative to training from scratch. This complementarity echoes findings from prior multimodal integration studies [23, 24], which demonstrated that combining chemical structure with phenotypic features improve bioactivity prediction, though those approaches were limited to closed-set settings with pre-computed features.

### Phenotypic imaging as a universal readout for biological queries

Our results suggest that Cell Painting images encode richer information about molecular activity than previously appreciated. The highest zero-shot accuracy on cell-based assays is expected given the direct correspondence with morphological readouts, but robust performance on bacterial assays despite training on mammalian cells suggests that phenotypic signatures capture conserved principles of chemical perturbation across organisms. Even biochemical assays achieved performance well above random, indicating that phenotypic profiles retain latent information about target-level activity. Cross-attention analysis shows that after training, the assay query network shifts attention from background regions to subcellular structures (Fig. 5c), providing suggestive evidence that the model grounds linguistic assay descriptions in visual phenotypes rather than imaging artifacts. These findings extend the “profile once, predict many” paradigm [9] from closed-set prediction to open-ended biological querying.

### Limitations and future directions

Performance on biochemical assays lags behind cell-based assays, consistent with the expectation that purified molecular interactions do not always produce distinct morphological changes [25]. We could not evaluate open-set performance on ChEMBL-209 because identical descriptions map to different activity thresholds, highlighting the need for standardized, machine-readable assay ontologies [26, 27]. Richer assay descriptions incorporating mechanistic hypotheses could further improve zero-shot transfer. Looking forward, integrating complementary modalities such as transcriptomics [3] or temporal imaging could capture dynamic responses that static snapshots miss. As assay ontologies mature, treating biological experiments as queryable questions rather than fixed classification targets may provide a scalable foundation for computational drug discovery. Prospective validation on genuinely novel assays will provide the strongest evidence for real-world utility.

## 4 Methods

In this paper, we aim to develop a model that predicts compound activity across diverse biological assays under three increasingly challenging generalization settings: (i) novel compounds within known assays, (ii) known compounds against unseen assays, and (iii) novel compounds against unseen assays.

We present **OpenPheno**, a multimodal framework that predicts whether a compound is active in a given assay, based on Cell Painting images of compound-treated cells, the compound’s chemical structure, and a natural-language description of the target assay. The framework employs a two-stage training paradigm: Stage I pre-trains a visual encoder on Cell Painting images via a combination of image-wise self-supervised learning and vision-smiles contrastive learning; Stage II introduces an assay query network (AQN) that conditions the prediction on natural-language assay descriptions, enabling zero-shot generalization to unseen assays. In the following sections, we will detail the problem formulation, model architecture, dataset construction, and experimental procedures.

### 4.1 Problem Formulation

We formalize bioactivity prediction over a set of compounds 𝒞 = {1, …, *N*} and biological assays 𝒜= {1, …, *M*}. Each compound *i* ∈ 𝒞 is associated with two modalities: a phenotypic image *x*_*i*_ (capturing morphological responses via Cell Painting) and a molecular structure *s*_*i*_ (encoded as a SMILES string [28]). Each assay *j* ∈ *𝒜* is semantically defined by a natural language description *t*_*j*_, which details its biological target and experimental protocol. We observe binary activity labels *y*_*ij*_ ∈ {0, 1} for a sparse subset of observed compound-assay pairs *𝒪*⊂ *𝒞* × *𝒜*.

#### Closed-set bioactivity prediction

Conventional approaches formulate bioactivity prediction as a closed-set multi-label classification problem:

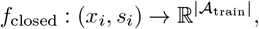

where 𝒜_train_ denotes a fixed set of assays observed during training. This formulation is inherently limited: it restricts inference exclusively to pre-defined biological endpoints and necessitates model retraining or architectural modifications to accommodate any new experimental targets.

#### Open-set bioactivity prediction

To overcome these limitations, we reframe bioactivity prediction as an open-set, visual-language question-answering (QA) task. In this paradigm, the assay description *t*_*j*_ acts as the biological “query” (the question), while the multimodal compound representation (*x*_*i*_, *s*_*i*_) serves as the universal evidence (the context). Formally, we learn a function:

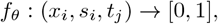

where *f*_*θ*_ estimates the probability that compound *i* is active in assay *j*. This formulation unlocks zero-shot generalization: at inference time, the model can predict activity for entirely novel assays 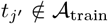 without any model retraining or bespoke biological experiments. By leveraging the semantic structure shared across natural language assay descriptions, the model effectively transfers knowledge to unseen experiments targeting novel biological mechanisms.

### 4.2 Framework Overview

As illustrated in Fig. 1, the proposed framework follows a two-stage training paradigm: we first pre-train a visual encoder on Cell Painting images using a dual objective that yields phenotypic representations that are both chemically grounded and invariant to batch effects; at stage two, we then cast bioactivity prediction as open-set question answering, with the Cell Painting images as visual evidence, chemical structures, and natural language assay descriptions as a query. This formulation enables zero-shot inference on assays not seen during training.

### 4.3 Stage I: Multimodal Representation Learning

We pre-train the visual encoder with a dual objective that enforces biological consistency and semantic alignment. This involves image-only self-supervised learning to ensure consistency across experimental replicates, and cross-modal contrastive learning to align phenotypic features with the molecule, denoted as the SMILES embedding. To suppress plate-level technical noise before cross-modal alignment, a gated fusion module explicitly contrasts compound-treated wells against DMSO controls from the same plate, retaining only treatment-specific phenotypic features for contrastive learning.

#### 4.3.1 Image-only Self-Supervised Learning

Biological replicates of the same compound across different plates should exhibit representational invariance; however, plate-specific batch effects-arising from variations in staining and imaging conditions-often obscure this correspondence. To enforce the cross-plate consistency, we adopt the training objective as proposed in DINO framework [29].

##### DINO

For a given compound, we sample a set of views *𝒱* = *𝒱*^*A*^∪*𝒱*^*B*^ derived from two replicate plates, *A* and *B*. Here, *𝒱*^*A*^ and *𝒱*^*B*^ contain both global crops (large field of view) and local crops (small field of view) from their respective plates. We train a student network to match the output of a momentum-updated teacher network across these plates:

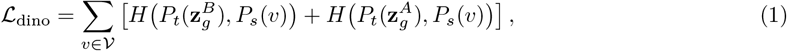

where *v* represents an individual image crop (view) sampled from the union set *𝒱*. 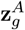 and 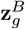 represent teacher embeddings of global crops from plates *A* and *B*, respectively. *P*_*t*_ and *P*_*s*_ denote the softmax distributions over the teacher and student outputs (with centering and sharpening applied to *P*_*t*_ to prevent collapse), and *H* is the cross-entropy loss. By using representations from one plate to predict the global context of a replicate plate, the encoder learns to suppress plate-specific artifacts while preserving compound-specific phenotypic signatures.

#### 4.3.2 Cross-modal Contrastive Learning

Phenotypic images and chemical structures offer complementary views of compound activity. By aligning these modalities, we establish a shared semantic manifold that grounds visual features in molecular structure, enabling knowledge transfer from large-scale chemical databases to phenotypically profiled compounds.

##### Gated phenotypic fusion

Cell Painting images frequently contain technical noise that can dominate the biological signal. To disentangle treatment effects from this background, we introduce a gated fusion module that explicitly contrasts treated wells against control wells from the same plate. Let *x*_*i*_ denote the image of a compound-treated well and 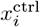 denote a DMSO-treated control well from the same plate. We extract the global image representations (i.e., the [CLS] token from the final layer) using the shared encoder:

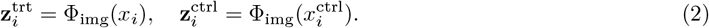

We combine these representations via a learnable gating mechanism to emphasize compound-specific deviations:

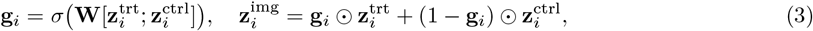

where [·;·] denotes concatenation, **W** is a linear projection mapping concatenated features to the gate dimension, *σ* is the sigmoid function, and ⊙denotes element-wise multiplication. The gate **g**_*i*_ effectively acts as a filter, retaining features modulated by the treatment while suppressing background noise shared with the control.

##### Vision-structure alignment

Chemical structures are encoded using a frozen MoleculeSTM encoder [30], yielding 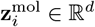. We project both the fused image representation 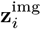 and the molecular representation 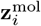 to a shared latent space. These projections are aligned via a symmetric InfoNCE loss [31]:

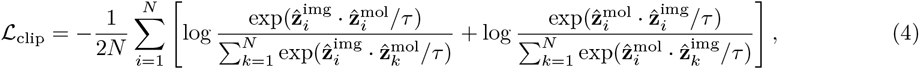

where 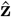 denotes the *L*_2_-normalized projection of the respective embedding, and *τ* is a learnable temperature parameter initialized to 0.07.

#### 4.3.3 Pretraining Objective

The visual encoder is pretrained by minimizing the weighted sum of the self-supervised and contrastive objectives:

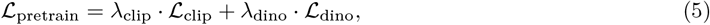

where *λ*_clip_ and *λ*_dino_ balance the contribution of each loss. Following pretraining, the vision encoder Φ_img_(·) is frozen for downstream zero-shot bioactivity prediction, and it is unfrozen for few-shot adaptation tasks. We also explored alternative objectives, including Masked Autoencoders (MAE) [18] and SigLIP [19]; however, both MAE and SigLIP underperformed relative to the standard InfoNCE loss (see Table S6 for detailed ablation results).

### 4.4 Stage II: Assay-Aware Bioactivity Prediction

At this stage, we introduce an assay query network (AQN) that implements the open-set bioactivity prediction. The AQN fuses the embeddings of chemical structure and assay description to construct a joint query, which then attends to the phenotypic image features to infer bioactivity. This design enables open-set generalization, *i*.*e*., the model can answer novel biological questions by interpreting new assay descriptions without retraining.

#### Image encoding

Given an input image *x*_*i*_, we extract the full sequence of embeddings, comprising both the global [CLS] token and the spatial patch tokens from the final layer of the pretrained visual encoder. This preserves both the global context and fine-grained spatial information:

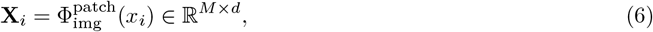

where *M* is the number of patches and *d* is the embedding dimension.

#### Chemical structure encoding

The molecular structure *s*_*i*_ (represented as a SMILES string) is encoded with a frozen MoleculeSTM encoder:

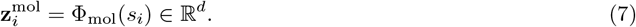

#### Assay description encoding

Raw assay metadata is first converted to language descriptions using GPT-5 [32], which extracts key biological entities (molecular target, organism, readout type) and crucial experimental variables (concentration, duration), for example, given the raw metadata {ASSAY_NAME: BAFComplex, ASSAY_DESC: esBAF, OBS_NAME: actin cp, ASSAY_TYPE: cell, READOUT_TYPE: qPCR}, the resulting description is: ‘This cell-based BAFComplex (esBAF) assay targets cytoplasmic actin (actin cp) in an unspecified organism, using qPCR to quantify gene expression. The goal is to assess esBAF-driven regulation of actin cp transcripts in cells’. Therefore, each assay *j* is associated with a language description *t*_*j*_ that specifies the biological question. These descriptions are then encoded using BioLord-2023 [17], a biomedical sentence embedding model optimized on UMLS and SNOMED CT ontologies [26].

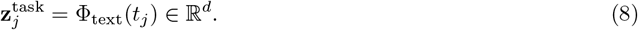

#### Assay query network (AQN)

The AQN constructs a dynamic query by fusing chemical structure and assay description embeddings through adaptive gating:

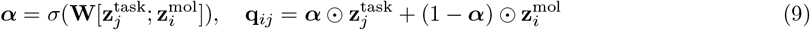

This allows the model to dynamically balance molecular identity versus biological context when forming the query. For some assays, chemical structure dominates; while for others, the assay descriptions are more informative. The learned gating mechanism automatically determines this balance based on the specific compound-assay pair.

The joint query **q**_*ij*_ then attends to the visual evidence (phenotypic image embeddings **X**_*i*_ corresponding to molecule *i*) via multi-head cross-attention:

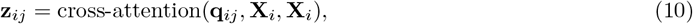

where the query **q**_*ij*_ selectively pools information from cellular regions in **X**_*i*_ relevant to answering the biological question. This cross-attention mechanism enables the model to focus on phenotypic features most informative for the specific assay being queried.

The attended representation **z**_*ij*_ is passed through an MLP classification head to predict the answer:

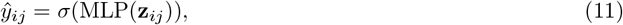

trained with binary cross-entropy loss over compound-assay pairs.

### 4.5 Datasets and Pre-processing

#### Phenotypic profiling data

All phenotypic images are derived from the Cell Painting assay [2], a high-content multiplexed microscopy protocol that captures morphological responses of cells to chemical perturbations. We evaluate our framework on two established benchmarks: **(i) Broad-270 [9]**: this benchmark comprises 16,170 compounds evaluated across 270 biological assays (63,089 compound–image pairs). Phenotypic images originate from Cell Painting screens conducted at the Broad Institute, while assay labels are derived from internal screening campaigns. **(ii) ChEMBL-209 [7]**: this benchmark comprises 10,574 compounds associated with 209 assays (47,304 compound-image pairs). While the phenotypic images are drawn from the same Broad Institute resource, bioactivity labels are aggregated from the ChEMBL database [33].

#### Image pre-processing

Raw Cell Painting images were processed following standard high-content imaging practices [2, 34]. Each image comprises five fluorescence channels capturing nuclei, endoplasmic reticulum, nucleoli and cytoplasmic RNA, actin and Golgi, and mitochondria. For each fluorescence channel, intensity values were clipped to the 0.1^th^ and 99.9^th^ percentiles to suppress hot pixels and extreme artifacts, followed by channel-wise min–max normalization. Following prior work [9], raw images consist of U2OS human osteosarcoma cells [35] imaged in 384-well plates. For each compound, six fields of view are stitched into a single montage of 1040× 2088 pixels. These montages are subsequently resized to 780× 1566 pixels while preserving the original aspect ratio. This resolution preserves subcellular morphological details while maintaining sufficient population context for robust phenotypic characterization. During the multimodal representation learning stage, we applied multi-scale image inputs, including global crops of size 224× 224 and local crops of size 96 ×96, to encourage scale-invariant feature learning. For downstream visual question answering, all images were standardized to 224× 224 pixels using random cropping during training and center cropping during validation and testing. This preprocessing strategy ensures consistent input dimensions while introducing controlled variability during training to improve generalization.

#### Assay curation and description construction

We constructed binary activity labels and assay descriptions following protocols established in the original studies.

- **Broad-270**. Assays were selected from a historical collection of screens performed at the Broad Institute. Redundant assays were removed based on Jaccard similarity of hit sets (threshold= 0.7), followed by hierarchical clustering with cosine distance. Continuous assay readouts were binarized using a double-sigmoid normalization [11], which fits separate sigmoid functions to positive and negative deviations from the median independently, thereby accommodating asymmetric response distributions. Normalized scores were then converted to binary activity labels via statistical thresholding, following the protocol established in [9]. To generate assay descriptions, raw metadata fields (assay name, description, observation name, and readout type) were aggregated into structured narratives using GPT-5. This process standardized terminology and extracted key biological entities. The prompt template are provided in Supplementary Methods.
- **ChEMBL-209**. Assays were aggregated from the ChEMBL database following prior work [7]. Compounds were labeled as active if they satisfied either: (i) a pChEMBL value exceeding 5.5, 6.5, or 7.5, or (ii) an explicit “Active” annotation. Consequently, a single biological assay often corresponds to multiple binary prediction targets reflecting different activity thresholds. However, the original ChEMBL metadata lacks the granularity to distinguish which threshold defines a specific label column. This ambiguity causes identical textual descriptions to map to distinct classification tasks, rendering the descriptions semantically insufficient for open-set querying. We therefore reserve ChEMBL-209 exclusively for closed-set evaluation.

All descriptions were standardized into concise, structured narratives emphasizing: (i) explicit identification of biological entities (molecular target, assay modality, cell line, organism, and readout type), and (ii) preservation of critical experimental variables such as compound concentration and exposure duration.

#### Assay-based dataset splitting

To ensure the held-out set represents the broader experimental landscape, we employed a stratified splitting strategy for the Broad-270 benchmark. We balanced the partition across three dimensions (Table S7): (i) biological assay category (*e*.*g*., cell-based, biochemical, bacterial), ensuring coverage of diverse mechanisms; (ii) assay sample size, preventing data-rich assays from dominating any single data split; and (iii) positive activity rate, ensuring comparable class imbalance across splits. This stratification guarantees that performance on the unseen test set reflects true generalization capability to novel biological questions. Note that ChEMBL-209 was excluded from this zero-shot setting due to semantic overlaps in its assay descriptions, and was reserved for standard closed-set evaluation.

#### Dataset construction for few-shot adaptation

To assess the model’s ability to adapt to new experimental contexts with minimal data, we constructed few-shot learning splits derived from the **54 unseen test assays**. For each assay, we further partitioned the available data into train, validation, and test sets. Crucially, this partition was conducted based on molecular scaffolds rather than random splitting, ensuring that compounds in the **test set** are structurally distinct from those in the **training set**, rigorously testing the model’s ability to generalize to novel chemical space rather than memorizing specific structures. We maintained the original ratio of sample sizes across assays to preserve the data distribution. Furthermore, we enforced a constraint that the final test set for each assay must contain at least one active and one inactive compound to enable valid metric calculation. We evaluated adaptation at labeled data fractions of 0.1%, 1%, 10%, and 100% of the available training data for each held-out assay. This setup simulates a realistic scenario in drug discovery, where a new assay is introduced with a limited set of screened diverse compounds, and the model must predict activity for structurally novel candidates.

### 4.6 Implementation

#### Stage-I: pretraining

We utilized a Vision Transformer (ViT-Base/16) [36] modified to accept 5-channel inputs as the vision encoder. The model was pre-trained for 100 epochs using the AdamW optimizer [37] with a batch size of 512. We employed a cosine learning rate schedule, starting from a base learning rate of 00201 ×10^*−*4^ (linearly scaled with batch size) and decaying to 1× 10^*−*6^, with a 10-epoch linear warmup. The weight decay followed a cosine schedule increasing from 0.04 to 0.4. The training objective combined CLIP and DINO losses with equal weighting (*λ*_clip_ = *λ*_dino_ = 1.0). For the DINO objective, the teacher network was updated via an exponential moving average (EMA) with the momentum parameter increasing from 0.996 to 1.0 during training. The teacher temperature was set to 0.04, while the student temperature was fixed at 0.1.

#### Stage-II: bioactivity prediction

We adapted the pre-trained vision backbone for multi-label bioactivity prediction using an assay query network (AQN) via a 4-layer Transformer Decoder (8 attention heads, 2048 feed-forward dimension, and 0.1 dropout). The model was trained using the AdamW optimizer with a learning rate of 3× 10^*−*4^ and a weight decay of 1× 10^*−*4^. The batch size was set to 512. We utilized a Masked Binary Cross-Entropy (BCE) loss to handle sparse labels in the dataset. To prevent overfitting, a dropout rate of 0.1 was applied across the AQN and projection layers, and gradients were clipped at a norm of 3.0. The model was trained for a maximum of 20 epochs, employing an early stopping strategy where the optimal checkpoint was selected based on the minimum validation loss.

#### Few-shot experiment

For the few-shot adaptation tasks, we initialized the model with the weights from the Stage II pre-trained assay query network (AQN). Unlike the zero-shot setting where the entire architecture is frozen, we unfroze the vision backbone. To prevent catastrophic forgetting of the pre-trained knowledge, we employed a lower learning rate of 1× 10^*−*6^. The model was fine-tuned using the AdamW optimizer with a batch size of 512. Training was conducted on the predefined data fractions (0.1%, 1%, 10%, and 100%) detailed in Section 4.5. The model was fine-tuned for up to 20 epochs. Consistent with Stage II, early stopping was applied by monitoring the validation loss, selecting the checkpoint with the lowest validation loss to prevent overfitting on the limited support sets.

### 4.7 Baselines

We compared OpenPheno against established baselines spanning two categories, evaluated under the standard compound-split setting where all assays are observed during training.

#### Supervised models

On ChEMBL-209, we compared against six convolutional architectures trained in a multi-task setting on raw microscopy images: ResNet-50 [13], DenseNet-121 [14], GapNet [7] (a custom architecture using global average pooling for high-resolution images), MIL-Net [15] (multiple instance learning), M-CNN [16] (multi-scale CNN), and SC-CNN [7]. We report original published results [7]. On Broad-270, we retrained ResNet-50, the best-performing architecture in this family, under identical data splits for fair comparison, as the remaining architectures were evaluated only on ChEMBL-209 in the original study.

#### Linear probing

This category encompasses feature-based approaches and contrastive pretraining methods. *Feature-based approaches*: on Broad-270, LateFusion [9] trains separate multi-task feedforward networks on pre-computed chemical structure (CS), Cell Painting morphology (MO), and gene expression (GE) features (approximately 1,700 CellProfiler features per well for morphology), before aggregating prediction scores post hoc. All pairwise and three-way modality combinations were evaluated; results are taken from the original publication as source code is not publicly available. On ChEMBL-209, we evaluated a feedforward network trained on CellProfiler morphological features [6, 38]. *Contrastive pretraining*: CLOOME [10] pre-trains a visual encoder via contrastive learning to align microscopy images with chemical structures, followed by linear probing for bioactivity prediction. We additionally report linear probes on our own frozen encoders (Vision Encoder and SMILES Encoder from Stage I pretraining) to isolate the contribution of the Assay Query Network.

All baselines above are architecturally limited to closed-set prediction, treating assays as fixed output indices. They cannot interpret novel assay descriptions or generalize to assays absent from training without complete retraining.

### 4.8 Evaluation Metrics

To evaluate predictive performance, we report the mean Area Under the Receiver Operating Characteristic curve (AUROC) and mean F1-score across all assays in the test set. We prioritize AUROC and F1-score over metrics such as Area Under the Precision-Recall Curve (AUPRC) or Enrichment Factor (EF), as the latter are highly sensitive to the underlying positive rate (class imbalance). Since the active rate varies significantly across different biological assays, AUPRC and EF can be misleading when aggregated.

AUROC was computed directly from predicted probabilities without thresholding. For F1-score, we determined the optimal decision threshold by maximizing the F1-score on the validation set, and applied this threshold to test set predictions. Model selection was based on the checkpoint achieving the lowest validation loss.

To estimate uncertainty, we computed 95% confidence intervals via bootstrapping: for each evaluation, we resampled the set of assays with replacement 1,000 times, computed the macro-averaged AUROC for each bootstrap sample, and reported the 2.5th and 97.5th percentiles as the lower and upper bounds of the confidence interval.

## 5 Data Availability

The Cell Painting images used in this study were made available by Bray et al. [39], and can be obtained from the following link: http://gigadb.org/dataset/100351. The Broad-270 [9] assay labels and compound metadata are available at https://github.com/CaicedoLab/2023_Moshkov_NatComm. The ChEMBL-209 benchmark data were obtained following the protocol described in Hofmarcher et al. [7] available at https://github.com/ml-jku/hti-cnn, with bioactivity labels derived from the ChEMBL database [33]. The processed datasets, including assay descriptions, data splits for all three evaluation settings, and few-shot adaptation partitions, will be made publicly available at https://github.com/Yuze-e20/OpenPheno

## 6 Code Availability

Code is publicly accessible at https://github.com/Yuze-e20/OpenPheno

## A Supplementary Materials

### A.1 Detailed experimental results

**Table S1.** Closed-set bioactivity prediction on Broad-270 (16,170 compounds, 270 assays). Late fusion baselines combining chemical structures (CS), morphology (MO), and gene expression (GE) are from Moshkov et al. All methods follow the standard compound-split protocol. Mean AUROC and counts of assays exceeding AUROC thresholds of 0.9 and 0.7 are reported.

| Type | Method | Mean AUROC | AUROC>0.9 | AUROC>0.7 |
| --- | --- | --- | --- | --- |
| Supervised | ResNet | 0.61 | 15 | 70 |
| Linear probing | Independent Modality(CS) | 0.63 | 22 | 88 |
| Linear probing | Independent Modality(MO) | 0.64 | 28 | 83 |
| Linear probing | Independent Modality(GE) | 0.59 | 22 | 57 |
| Linear probing | LateFusion(CS+MO) | 0.66 | 29 | 99 |
| Linear probing | LateFusion(GE+MO) | 0.65 | 29 | 86 |
| Linear probing | LateFusion(CS+GE) | 0.65 | 24 | 87 |
| Linear probing | LateFusion(CS+MO+GE) | 0.67 | 28 | 96 |
| Linear probing | Vision Encoder(CLIP+DINO) | 0.65 | 23 | 89 |
| Linear probing | SMILES Encoder(CLIP+DINO) | 0.63 | 22 | 84 |
| Linear probing | SMILES Encoder(CLIP+DINO) | 0.64 | 15 | 78 |
| AQN | OpenPheno | **0.71** | **32** | **106** |

**Table S2.** Closed-set bioactivity prediction on ChEMBL-209 (10,574 compounds, 209 assays). All methods follow the standard compound-split protocol with scaffold-based partitioning. Supervised baselines are from Hofmarcher et al. Mean AUROC and counts of assays exceeding AUROC thresholds of 0.9 and 0.7 are reported.

| Type | Method | Mean AUROC | AUROC>0.9 | AUROC>0.7 |
| --- | --- | --- | --- | --- |
| Supervised | ResNet | 0.73 | 68 | 119 |
| Supervised | DenseNet-121 | 0.73 | 61 | 121 |
| Supervised | GapNet | 0.73 | 63 | 117 |
| Supervised | MIL-Net | 0.71 | 61 | 105 |
| Supervised | M-CNN | 0.71 | 57 | 105 |
| Supervised | SC-CNN | 0.71 | 61 | 109 |
| Supervised | FNN | 0.68 | 55 | 90 |
| Linear probing | CLOOME | 0.71 | 57 | 109 |
| Linear probing | CellProfiler | 0.66 | 35 | 84 |
| Linear probing | Vision Encoder (CLIP+DINO) | 0.72 | 61 | 109 |
| Linear probing | SMILES Encoder (CLIP+DINO) | 0.67 | 2 | 112 |
| AQN | OpenPheno | **0.76** | 68 | 137 |

**Table S3.** Zero-shot bioactivity prediction performance of OpenPheno on 54 unseen assays from Broad-270. Mean AUROC and F1 score are reported for each biological assay category. No labeled examples were provided for any test assay, so closed-set baselines cannot operate in this setting. 95% confidence intervals are reported.

| Metric | Overall | Cell | Biochem | Bacterial | Fungal & Yeast |
| --- | --- | --- | --- | --- | --- |
| AUROC | 0.75 (0.70, 0.80) | 0.79 (0.74, 0.83) | 0.65 (0.61, 0.70) | 0.73 (0.68, 0.78) | 0.79 (0.74, 0.83) |
| F1 | 0.25 (0.21, 0.29) | 0.31 (0.27, 0.35) | 0.15 (0.12, 0.17) | 0.11 (0.03, 0.18) | 0.28 (0.23, 0.33) |

**Table S4.** Few-shot adaptation performance across assay categories and data fractions on 54 unseen assays from Broad-270. OpenPheno is compared against SMILES Encoder, Vision Encoder (single-modality linear probes), and Scratch (AQN without pretraining) under data fractions from 0% (zero-shot) to 100% (full supervision). Mean AUROC with 95% confidence intervals are reported.

| Model | 0% | 0.1% | 1% | 10% | 100% |
| --- | --- | --- | --- | --- | --- |
| *Overall* |  |  |  |  |  |
| SMILES Encoder | – | 0.52 (0.43, 0.61) | 0.57 (0.48, 0.67) | 0.61 (0.52, 0.69) | 0.63 (0.55, 0.72) |
| Vision Encoder | – | 0.45 (0.35, 0.56) | 0.47 (0.36, 0.58) | 0.57 (0.46, 0.67) | 0.64 (0.53, 0.74) |
| Scratch | 0.69 (0.61, 0.77) | 0.68 (0.60, 0.76) | 0.69 (0.61, 0.77) | 0.69 (0.61, 0.77) | 0.72 (0.67, 0.77) |
| OpenPheno | **0.73** (0.65, 0.80) | **0.74** (0.66, 0.81) | **0.74** (0.66, 0.81) | **0.75** (0.68, 0.82) | **0.75** (0.67, 0.83) |
| *Cell* |  |  |  |  |  |
| SMILES Encoder | – | 0.54 (0.45, 0.63) | 0.58 (0.48, 0.68) | 0.64 (0.56, 0.73) | 0.64 (0.55, 0.73) |
| Vision Encoder | – | 0.43 (0.33, 0.55) | 0.43 (0.33, 0.55) | 0.59 (0.48, 0.70) | 0.69 (0.59, 0.79) |
| Scratch | 0.75 (0.66, 0.82) | 0.74 (0.66, 0.82) | 0.75 (0.66, 0.82) | 0.76 (0.68, 0.83) | 0.76 (0.69, 0.82) |
| OpenPheno | **0.77** (0.69, 0.84) | **0.77** (0.70, 0.84) | **0.78** (0.70, 0.84) | **0.79** (0.71, 0.85) | **0.79** (0.71, 0.84) |
| *Biochem* |  |  |  |  |  |
| SMILES Encoder | – | 0.41 (0.31, 0.50) | 0.57 (0.47, 0.65) | 0.57 (0.48, 0.65) | 0.59 (0.51, 0.65) |
| Vision Encoder | – | 0.41 (0.31, 0.51) | 0.48 (0.38, 0.57) | 0.52 (0.42, 0.62) | 0.57 (0.47, 0.66) |
| Scratch | **0.63** (0.540, 0.711) | 0.61 (0.52, 0.70) | 0.62 (0.53, 0.70) | 0.62 (0.52, 0.71) | 0.66 (0.55, 0.75) |
| OpenPheno | 0.62 (0.54, 0.69) | **0.63** (0.54, 0.71) | **0.63** (0.53, 0.71) | **0.64** (0.54, 0.72) | **0.67** (0.57, 0.76) |
| *Bacterial* |  |  |  |  |  |
| SMILES Encoder | – | 0.54 (0.47, 0.61) | 0.56 (0.46, 0.63) | 0.51 (0.47, 0.55) | 0.59 (0.49, 0.67) |
| Vision Encoder | – | 0.56 (0.44, 0.66) | 0.65 (0.54, 0.74) | 0.60 (0.47, 0.70) | 0.56 (0.44, 0.66) |
| Scratch | 0.59 (0.50, 0.67) | 0.61 (0.53, 0.68) | 0.60 (0.51, 0.68) | 0.62 (0.53, 0.69) | **0.74** (0.47, 1.01) |
| OpenPheno | **0.71** (0.63, 0.78) | **0.72** (0.64, 0.79) | **0.72** (0.63, 0.78) | **0.71** (0.62, 0.78) | 0.73 (0.63, 0.81) |
| *Fungal & Yeast* |  |  |  |  |  |
| SMILES Encoder | – | 0.71 (0.62, 0.78) | 0.58 (0.50, 0.66) | 0.54 (0.42, 0.64) | 0.78 (0.70, 0.83) |
| Vision Encoder | – | 0.59 (0.45, 0.72) | 0.50 (0.37, 0.61) | 0.48 (0.35, 0.59) | 0.49 (0.37, 0.60) |
| Scratch | 0.58 (0.48, 0.67) | 0.52 (0.43, 0.61) | 0.57 (0.47, 0.66) | 0.52 (0.42, 0.61) | 0.65 (0.46, 0.84) |
| OpenPheno | **0.79** (0.70, 0.87) | **0.80** (0.72, 0.87) | **0.81** (0.74, 0.88) | **0.81** (0.72, 0.88) | **0.80** (0.70, 0.88) |

**Table S5.** Comparison of Setting 2 and Setting 3. Performance comparison of zero-shot bioactivity prediction between Setting 2 (known compounds evaluated on novel assays) and Setting 3 (novel compounds evaluated on novel assays) across different biological assay categories. Mean AUROC and F1 scores along with their 95% confidence intervals are reported.

| Setting | Metric | Overall | Cell | Biochem | Bacterial | Fungal & Yeast |
| --- | --- | --- | --- | --- | --- | --- |
| Setting 2 | AUROC | 0.73 (0.65, 0.80) | 0.77 (0.70, 0.84) | 0.62 (0.54, 0.69) | 0.71 (0.63, 0.78) | 0.79 (0.70, 0.87) |
|  | F1 | 0.30 (0.24, 0.35) | 0.36 (0.31, 0.42) | 0.17 (0.15, 0.19) | 0.16 (0.06, 0.26) | 0.36 (0.28, 0.45) |
| Setting 3 | AUROC | 0.66 (0.57, 0.75) | 0.73 (0.64, 0.81) | 0.58 (0.49, 0.67) | 0.58 (0.49, 0.67) | 0.46 (0.35, 0.58) |
|  | F1 | 0.24 (0.21, 0.27) | 0.33 (0.28, 0.37) | 0.12 (0.10, 0.13) | 0.09 (0.09, 0.09) | 0.12 (0.12, 0.12) |

**Table S6.** Ablation study of pretraining strategies on zero-shot bioactivity prediction across assay categories. All models use the AQN architecture; only the Stage I pretraining objective differs. Pretraining variants include Scratch (no pretraining), MAE, CLIP, DINO, SigLIP, and their combinations. Mean AUROC with 95% confidence intervals on 54 unseen test assays are reported.

| Method | Overall | Cell | Biochem | Bacterial | Fungal & Yeast |
| --- | --- | --- | --- | --- | --- |
| Scratch | 0.72 (0.66, 0.76) | 0.75 (0.70, 0.80) | **0.67** (0.62, 0.71) | 0.63 (0.57, 0.67) | 0.70 (0.64, 0.76) |
| MAE | 0.70 (0.65, 0.74) | 0.73 (0.68, 0.78) | 0.66 (0.61, 0.69) | 0.64 (0.59, 0.68) | 0.66 (0.59, 0.72) |
| CLIP | 0.74 (0.68, 0.78) | 0.77 (0.72, 0.81) | 0.65 (0.60, 0.69) | 0.71 (0.65, 0.76) | 0.74 (0.67, 0.80) |
| DINO | 0.74 (0.69, 0.78) | **0.80** (0.75, 0.83) | 0.59 (0.55, 0.63) | 0.71 (0.65, 0.76) | **0.79** (0.74, 0.83) |
| SigLIP | 0.73 (0.67, 0.77) | 0.77 (0.72, 0.81) | 0.65 (0.60, 0.69) | 0.70 (0.63, 0.75) | 0.64 (0.56, 0.72) |
| SigLIP+DINO | 0.74 (0.69, 0.79) | 0.79 (0.73, 0.83) | 0.63 (0.58, 0.67) | 0.72 (0.65, 0.76) | 0.77 (0.71, 0.81) |
| CLIP+DINO | **0.75** (0.70, 0.79) | 0.79 (0.74, 0.83) | 0.65 (0.60, 0.69) | **0.73** (0.68, 0.78) | 0.79 (0.73, 0.83) |

**Table S7.** Statistical evaluation of the assay-wise dataset partition for the Broad-270 assay-split setting. Assays are stratified into training (189 assays, 70%), validation (27 assays, 10%), and test (54 assays, 20%) splits. The partition preserves comparable distributions of assay type, sample size, and positive rate across splits, ensuring that the test set performance reflects generalization to representative novel biological questions rather than a distribution shift.

|  | Train | Validation | Test |
| --- | --- | --- | --- |
| *Assay type distribution (%)* |  |  |  |
| Cell-based | 57.14 | 59.26 | 59.26 |
| Biochemical | 21.16 | 22.22 | 24.07 |
| Bacterial | 11.64 | 11.11 | 9.26 |
| Yeast | 6.88 | 7.41 | 5.56 |
| *Sample size per assay* |  |  |  |
| Mean | 2,367.9 | 1,666.9 | 1,720.2 |
| Median | 89.0 | 77.0 | 67.0 |
| *Positive rate (%)* |  |  |  |
| Mean | 8.45 | 6.08 | 8.66 |

### A.2 Prompt Sets

#### System Prompt for Broad-270

~~~
You are an expert biologist and data curator. Your task is to summarize
experimental metadata into a concise, scientifically accurate paragraph suitable for BioLord -2023 embedding generation.
Requirements :
1. Focus on Semantics: Explicitly mention the biological target, the organism, the method, and the goal.
2. Capture All Details: Meticulously include specific experimental conditions if present, such as duration (e. g., “ for 48 hours “), dosage, or specific timepoints. Do not omit these quantitative details even if they appear minor.
3. Conciseness: Keep it between 30 -60 words.
4. Output Format: You must output ONLY a valid JSON object. Do not include any conversational text.
JSON Structure :
{
  “ id “: “ The provided PUMA_ASSAY_ID “,
  “ summary “: “ The generated summary text”
}
~~~

#### User Prompt for Broad-270

~~~
- Here is the metadata for the assay to be summarized:
- Assay Name : { row [‘ ASSAY_NAME ‘]}
- Assay Description : { row [‘ ASSAY_DESC ‘]}
- Target/Strain : { row [‘ OBS_NAME ‘]}
- Assay Type : { row [‘ ASSAY_TYPE ‘]}
- Readout: { row [‘ READOUT_TYPE ‘]}
Please output the JSON object for ID: { row [‘ PUMA_ASSAY_ID ‘]}
~~~

#### System Prompt for ChEMBL-209

~~~
You are an expert biologist and data curator. Your task is to summarize experimental metadata into a concise, scientifically accurate paragraph suitable for BioLord -2023 embedding generation.
Requirements:
1. Focus on Semantics: Explicitly mention (if available) the biological target, the assay type, the organism, the method, the readout type and the goal.
2. Capture All Details: Meticulously include specific experimental conditions if present, such as duration (e. g., “ for 48 hours “), dosage, or specific timepoints. Do not omit these quantitative details even if they appear minor.
3. Conciseness: Keep it between 30 -60 words.
4. Output Format: You must output ONLY a valid JSON object. Do not include any conversational text.
JSON Structure:
{
  “ id “: “ The provided Unique_ID “,
  “ summary “: “ The generated summary text”
}
~~~

#### User Prompt for ChEMBL-209

~~~
Here is the metadata for the assay to be summarized:
-Assay Description: { row [‘ Description ‘]}
Please output the JSON object for ID: { row [‘ Unique_ID ‘]}
~~~

## Notes

https://github.com/Yuze-e20/OpenPheno

### Competing Interest Statement

The authors have declared no competing interest.

